# Adhesion force spectroscopy with nanostructured colloidal probes reveals nanotopography-dependent early mechanotransductive interactions at the cell membrane level

**DOI:** 10.1101/2020.01.02.892919

**Authors:** M. Chighizola, A. Previdi, T. Dini, C. Piazzoni, C. Lenardi, P. Milani, C. Schulte, A. Podestà

**Affiliations:** C.I.Ma.I.Na. and Dipartimento di Fisica “Aldo Pontremoli”, Università degli Studi di Milano, via Celoria 16, 20133 Milan, Italy

**Keywords:** Mechanosensing, mechanotransduction, mechanobiology, cell adhesion, nanotopography, nanostructured materials, cell microenvironment, extracellular matrix, atomic force microscopy, colloidal probes, adhesion force spectroscopy, integrin binding, adhesion complexes.

## Abstract

Mechanosensing, the ability of cells to perceive and interpret the microenvironmental biophysical cues (such as the nanotopography), impacts strongly on cellular behaviour through mechanotransductive processes and signalling. These events are predominantly mediated by integrins, the principal cellular adhesion receptors located at the cell/extracellular matrix (ECM) interface.

Because of the typical piconewton force range and nanometre length scale of mechanotransductive interactions, achieving a detailed understanding of the spatiotemporal dynamics occurring at the cell/microenvironment interface is challenging; sophisticated interdisciplinary methodologies are required. Moreover, an accurate control over the nanotopographical features of the microenvironment is essential, in order to systematically investigate and precisely assess the influence of the different nanotopographical motifs on the mechanotransductive process.

In this framework, we were able to study and quantify the impact of microenvironmental nanotopography on early cellular adhesion events by means of adhesion force spectroscopy based on innovative colloidal probes mimicking the nanotopography of natural ECMs.

These probes provided the opportunity to detect nanotopography-specific modulations of the molecular force loading dynamics and integrin clustering at the level of single binding events, in the critical time window of nascent adhesion formation. Following this approach, we found that the nanotopographical features are responsible for an excessive force loading in single adhesion sites after 20 – 60 s of interaction, causing a drop in the number of adhesion sites. However, by manganese treatment we demonstrated that the availability of activated integrins is a critical regulatory factor for these nanotopography-dependent dynamics.

## 1. INTRODUCTION

A complex crosstalk between cells and their microenvironment, i.e. the extracellular matrix (ECM), governs the development and maintenance of multicellular tissues. The biophysical properties of the microenvironment were therein identified as critical factors for the regulation of many cellular responses, such as proliferation, migration, and differentiation. The ECM is a complex meshwork of intertwined macromolecules (with protein and sugar components) characterised by the presence of fibrillary and reticular structures, pores and asperities at the nanoscale. The configurations can be relatively ordered, as e.g. in fibrillary collagen-dominated ECM, or instead rather disordered, as in basement membranes or brain ECM. However, on the local nanoscale level often irregularities, anisotropies and density gradients are present^1–6^.

In recent years, it became evident that mechanical stimuli of the ECM, such as rigidity and/or spatial organisation and dimensionality of adhesion sites (e.g., in terms of geometry and topography), alter intrinsic cellular properties, such as the actin cytoskeletal organisation/mechanics and the signalling status^7–10^. The intricate processes through which the cell perceives and reacts to mechanical and structural stimuli in its microenvironment were termed mechanosensing and mechanotransduction^11–18^.

Mechanotransductive processes are involved in virtually all aspects of the cellular life and tissue organisation^11–19^ and aberrations in components that participate to mechanosensing and mechanotransduction have been linked to various diseases, in particular in cancer and metastasis. A detailed comprehension of how biophysical ECM characteristics modulate mechanotransduction would promote new approaches for treatments of diseases, drug therapies or diagnostic approaches, exploiting and targeting identified mechanotransductive key regulators or structures^11, 20–25^.

Cells sense the biophysical microenvironmental information at the nanoscale, and the nanotopography emerged as a crucial parameter in the regulation of mechanotransductive processes and signalling^7–9, 26–29^. The mechanotransductive pathway is primarily mediated by specific transmembrane proteins, called integrins, and modulated by the extent to which these integrins cluster together into integrin adhesion complexes (IACs) and mature into bigger structures, such as focal complexes and focal adhesions (FAs). The extent of integrin clustering and FA maturation, in turn, depends on the force loading within the so-called molecular clutches of nascent adhesions, i.e. the initial connection of the ECM-binding integrins to forces generated by the actin cytoskeleton *via* adaptor proteins (in particular talin and vinculin). The spatial organisation of integrin adhesion sites (ligands) exerts a fundamental influence on the integrin clustering and eventually on cellular behaviour and responses. These effects are mediated in particular through the principal elements of mechanotransductive signalling, such as RhoGTPases, the actin cytoskeleton and mechanosensitive transcription factors (e.g., YAP)^11–19^.

However, a better understanding and quantification of the dynamics in the cell/microenvironment interface and the force development in the early steps of cellular mechanosensing in response to different topographical stimuli is required. In particular, although the presence of nanotopographical disorder (due to the impact on the spatial distribution of integrin ligands) has been shown to have a strong influence on protein adsorption, cell adhesion, integrin clustering and differentiation^7, 30, 31^, the systematic characterisation of the influence of disordered configurations with different nanoscale three-dimensional features is still in its infancy.

To unravel the molecular mechanisms of cellular mechanosensing and the mechanotransductive responses it provokes in the cells, versatile biophysical approaches are essential^10, 17, 29, 32–39^. Smart biomaterials with complex structural architectures and/or tuneable physical properties are needed to create controllable cellular microenvironments that mimic the *in vivo* ECM situation^7, 29, 32, 34, 35^. In addition, sophisticated techniques are required that enable the mapping of mechanobiologically relevant alterations in cells^17, 33, 36^.

In this context, atomic force microscopy (AFM) represents a powerful tool due to its capacity to allow an accurate probing of cell surfaces, determining cellular mechanical properties^40–43^ (and mechanotransduction-related alterations^44–46)^, and measuring adhesion forces down to the single-molecule contribution^36^. The standard techniques used to test the cellular adhesion properties are Single-Molecule Force Spectroscopy (SMFS)^47^ and Single-Cell Force Spectroscopy (SCFS)^48, 49^. SMFS consists in a functionalisation of the AFM probe (e.g., coating with ECM proteins) in order to permit the interaction with specific transmembrane proteins. SCFS instead uses the cell itself as a probe (attached to the cantilever) that interacts with the substrates of interest (including other cells).

SCFS has been widely used to study in detail the cooperative action of integrins in early cellular adhesion to fibronectin or collagen^50–52^ and their connection to internal cellular biochemical signalling^53^. This technique has furthermore been exploited to test the biocompatibility of materials for implants in orthopaedic surgery^54^, or the role of ligand spacing in the cell adhesion using substrates decorated with suitably functionalised nanoparticles, separating integrin ligands with different specific distances^55, 56^. A limitation of SCFS, as explained in detail by Naganuma^57^, is that the actual contact area between the cell and the substrates cannot be measured. As a matter of fact, the cell, once captured and immobilised on the tipless cantilever, will evolve its own adhesion on the probe itself, changing its morphology during time and with it the adhesion properties. These dynamics will introduce a bias in the force spectroscopy experiment and, since different cells behave differently, make results less comparable between each other.

To address these issues, we propose a reversal of the conventional SCFS approach. Our novel strategy for the study of early mechanotransductive interactions at the cell membrane level is based on the use of functionalised colloidal probes (CPs) mimicking the peculiar nanoscale topographical features of *in vivo* ECM for force spectroscopy experiments on living cells. The probes represent the source for the mechanical cues regulating the cascade of the mechanotransductive events. By inverting the typical cell-microenvironment interaction geometry (Figure 1), we take in particular control over the cell/substrate contact area, and obtain a more accurate assessment of the forces and molecular interactions that develop at the cell membrane during the early mechanotransductive events.

**Figure 1:**
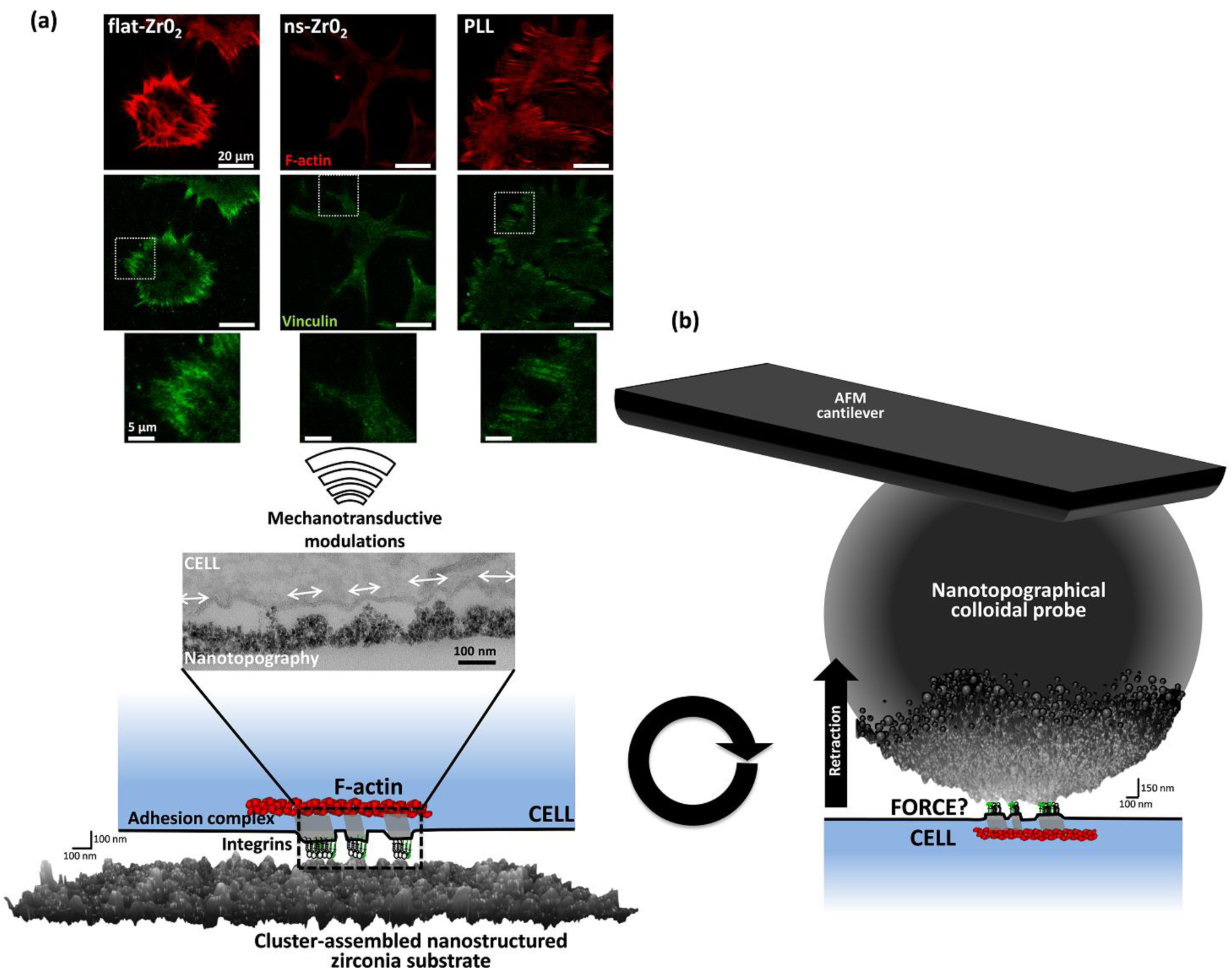
**(a)** The scheme at the bottom left illustrates the conventional approach used to study the impact of nanotopographical features on mechanotransductive processes. The image inset shows a transmission electron microscopy (TEM) image of an interaction sites between a PC12 cell and the asperities of cluster-assembled ECM-mimicking nanotopographical zirconia substrate (indicated by the white arrows) with an rms roughness r_q_ = 15 nm (ns-ZrO_2_), for which we have shown that it induces mechanotransductive modulations^44^. The immunofluorescence (IF) images in the panel above (in green, total internal reflection recordings of vinculin staining; in red, epifluorescence recordings of the actin filaments marked by phalloidin staining) demonstrate that, after 4 hours of interaction on the nanotopographical ns-ZrO_2_ substrate, only small punctate focal complex size adhesion sites formed and stress fibres were absent, whereas on the flat zirconia (flat-ZrO_2_) and PLL-coated glass (PLL) substrates, focal adhesions and stress fibres are present^44^. **(b)** Schematic illustration of the approach to measure adhesion forces to colloidal probes with a nanotopographical surface (r_q_ = 15 nm) produced by SCBD (the scale bars close to the nanostructured surfaces refer only to the nanotopography, other graphical icons are symbolic and not in scale). TEM and IF images in **(a)** were adapted from Schulte et al.^44^.

Our technological approach for the fabrication of nanotopographical surfaces consists in growing a cluster-assembled thin film of nanostructured zirconium oxide (ns-ZrO_2_, zirconia) with disordered, yet controlled, topographical features by means of supersonic cluster beam deposition (SCBD)^58–60^. Zirconia is a biocompatible material used in various clinical application, particularly for its chemical inertness and structural properties^61^. SCBD is an additive technique and the disordered surface morphology of the deposited films is characterised by nanoscale roughness and other morphological parameters, like the surface area or the correlation length, which can be accurately tuned and reproduced by controlling the film thickness through the deposition time^62^.

The obtained disordered nanostructured films possess nanotopographical features that resemble those that can be observed in *in vivo* ECM (e.g. in basement membranes or in the brain^5, 6^) and with nano-3D configurations (in terms of asperity dimensionality and distances) that have a potential to modulate integrin-dependent mechanotransductive processes and signalling^30, 44, 58, 63–67^ (Figure 1a). Indeed, also in those ECMs whose structure is dominated by fibrillar features, there are often irregularities at the nanoscale, e.g. in regards to distances and heights of fibres or in the pore sizes of reticular, crosslinked structures, which are well represented by the nanotopographical features of the ns-ZrO_2_^6, 44^.

Using these nanostructured thin films as substrates (Figure 1a), we have recently shown that the interaction of cells, in particular neuronal cells, with the nanotopographical features, impacts decisively on mechanotransductively relevant events, such as FA maturation, cytoskeletal organisation/mechanics and integrin signalling, as well as the cellular program and differentiative behaviour^44, 58, 63, 64, 68^.

Instead of plating the cells on the nanostructured substrates, here we exploited the nanotopographical colloidal probes (nt-CPs) to stimulate the cells and to characterise the interfacial integrin dynamics by means of adhesion force spectroscopy (Figure 1b). Compared to our previous experiments, this approach provided access to phenomena that occur at the cell-microenvironment interface that would otherwise be buried below the cell body.

To date, only few protocols have been developed to functionalise AFM probe surfaces either with ECM proteins, like collagens, laminins, or cellulose nanofibers^69, 70^ or with nanoparticles^71–73^. In the former case, natural biomaterials were typically deposited onto CPs, but poor or null control over the nanoscale topography was possible. In the latter case, AFM standard sharp tips were decorated with nanoparticles in order to study their interaction with cells; in other experiments the AFM tip apex was used to model a single nanoparticle^74^. Our nt-CPs were instead used to study the early steps of the cell adhesion to an ECM-mimicking nanotopography with the sensitivity of adhesion force spectroscopy measurements. Following this innovative approach, we were able to follow the temporal evolution of early integrin-related adhesion events at the interface between the cell and a nanotopographical microenvironment at the level of single adhesion events and piconewton force range. To our best knowledge, in this framework of cell/nanotopography interaction such a resolution has not been achieved to date.

## 2. MATERIALS AND METHODS

### 2.1 Fabrication and calibration of the colloidal probes

#### Fabrication of the colloidal probe

The procedure for the fabrication of colloidal probes is based on the approach described in detail in Ref.^75^. Borosilicate glass spheres (Thermo Fisher Scientific), with radius R = 10 ± 1μm, are first cleaned to remove surface contaminants. The cleaning procedure consists in three sequential 60 seconds centrifugations (10.000 rpm) in a 1:1 water and ethanol solutions, carefully replacing the old with new solution after each centrifugation. The cleaned spheres are then dispersed in toluene and deposited on a microscopy glass slide coated with a thin Au film (with thickness 100 nm) deposited by sputtering. The Au film is used to reduce the capillary force between the sphere and the substrates with respect to that between the sphere and a tipless cantilever^75^ (Micromash HQ:CSC38/tipless/no Al, force constant *k* = 0.02-0.03 N/m). The capture of the sphere by the cantilever is done using the XY motorised stage of the AFM microscope, integrated in the optical inverted microscope. The probe is then transferred in a pre-heated high-temperature oven and kept for 2 hours at 780°C. This temperature is slightly below the softening point of borosilicate glass, which is qualitatively defined as the temperature at which a solid object begins collapsing under its own weight. After two hours, the microsphere is covalently attached to the cantilever. Due to the monolithic character of the resulting CP, any gold residue, as well as other contaminants, can be effectively removed by washing the probe in aqua regia (a mixture of nitric and hydrochloric acids, in a molar ratio of 1:3), or any other aggressive solution.

#### Determination of the probe radius

The characterisation of the CP radius is performed by AFM reverse imaging of the probe on a spiked grating (TGT1, Tips Nano), as detailed in Ref.^75^. Upon scanning the CP on the spiked grating, hundreds of independent replicas of the probe apical region are obtained. From the measured geometrical properties (like the volume V and the height h) it is possible to determine the value of the radius R by fitting a spherical cap model *V* = *π*/3*h*^2^(3*R* − *h*) to the data. The evaluated probe radius has an accuracy as good as 1%.

#### Calibration of the cantilever spring constant

The spring constant is calibrated using the thermal noise method^76, 77^ where a special correction factor is applied in order to take into account the relevant dimension and mass of the glass sphere^78^. For large CPs, the conditions that both dimension and mass of the sphere are small compared to length and mass of the cantilever are not always satisfied. Since the mass of the microsphere scales with the cube of the radius *R*, beads with radii larger than 5 μm possess a mass comparable to the mass of the cantilever, and in some cases, especially with stiffer, shorter cantilevers, even larger. These conditions lead to the failure of the assumption that the mass of the cantilever is uniformly distributed along its length, resulting in an underestimation of the spring constant.

According to Ref.^78^, it is possible to correct the apparent spring constant *K*^t*h*^ measured by the thermal noise method using the formula:

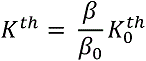

where *K*^t*h*^ is the corrected spring constant, while for rectangular cantilevers *β_0_* = 0.817. *β* depends on the reduced mass of the sphere *m*∽, i.e. the ratio of the mass of the sphere *m_S_* to the mass of the cantilever *m_C_*, and on the reduced gyration radius of the sphere *r*∽, proportional to the ratio of the radius *R* of the sphere to the length *L* of the cantilever. For small tips, *m*∽ and *r*∽ are negligibly small, and *β*= *β*_0_, therefore *K*^t*h*^ = *K*^t*h*^. In our case, since *L* = 350 μm, *m_C_* = 2.65×10^3^ ng, *m_S_* = 1.05×10^3^ ng, we had *m*∽ = 0.4 and *r*∽ = 0.03, therefore corrections were needed.

### 2.2 Production of nanotopographical and reference CPs

#### Deposition of the ns-ZrO_2_ film on the CP

For the production of nt-CPs, ns-ZrO_2_ films are deposited on the colloidal probes exploiting an SCBD apparatus equipped with a Pulsed Microplasma Cluster Source^79, 80^ (Figure 2). Partially oxidised zirconia clusters are produced within the PMCS and then extracted into the vacuum through a nozzle to form a seeded supersonic beam. Clusters are collected directly on the CPs intercepting the beam, in the deposition chamber. Upon landing on the probe surface, which is locally flat due to the large radius, clusters form a nanostructured, highly porous, high-specific area, biocompatible ns-ZrO_2_ film^44, 58, 59, 81, 82^. The oxidation of the nanostructured film further proceeds upon exposure to air, up to an almost complete stoichiometry, although rich of local defects. The crystalline phase is cubic at room temperature^62^.

**Figure 2:**
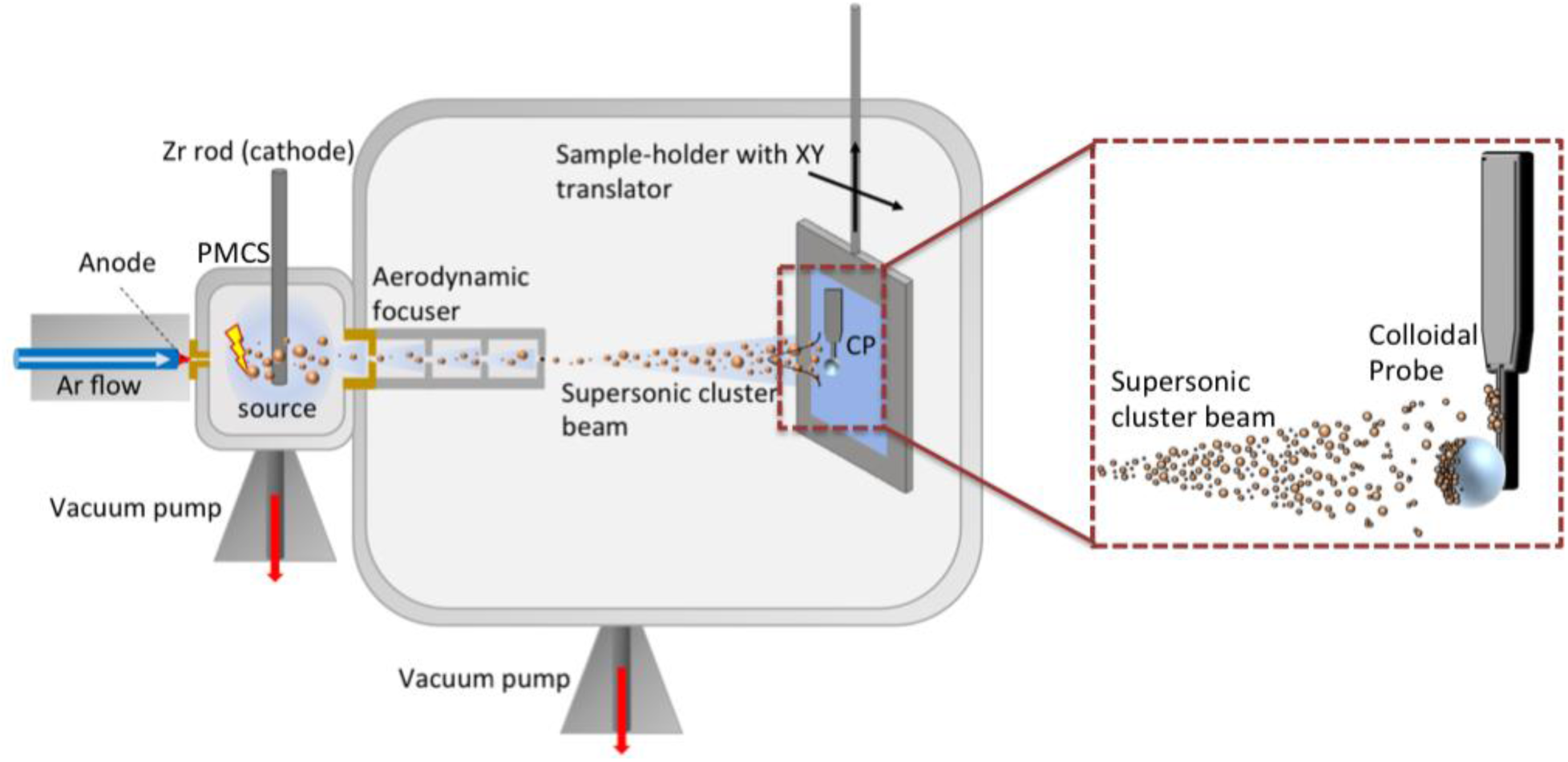
A schematic representation of the SCBD apparatus for the production of nt-CPs. The Zr rod is sputtered by a plasma discharge, triggered by the introduction of argon (Ar) through a pulsed valve into the source cavity and the application of a high voltage between the Zr rod and the anode. The ablated species condense into clusters and the resulting gas-clusters mixture is extracted through a nozzle and an aerodynamic focuser into a high vacuum chamber; during the process, the carrier gas-clusters mixture undergoes a supersonic expansion. The cluster beam impinges on the CP, where a thin, nanostructured ZrO_2_ film is formed.

The root mean square (rms) roughness *r_q_* of the deposited film, defined as the standard deviation of surface height values, evolves with the film thickness *h*, according to the power law^59, 60^ *r*_*q*_∼ *h*^*b*^, where *b* = 0.35 approximately (details about the evolution of the roughness-thickness relation and on the mechanical stability of the nanostructured thin film can be found in the Supporting Information, Figures S1,2)^60, 62^. Therefore, by controlling the thickness of the film deposited using a quartz microbalance placed inside the deposition chamber, it is possible to produce nanotopographical CPs with very high reproducibility. The typical thickness of the ns-ZrO_2_ films deposited on CPs was in the 70-250 nm range, corresponding to roughness values in the range 13-20 nm.

#### Deposition of smooth ZrO_2_ films on the CP

Thin, compact and very smooth coatings of ZrO_2_ (flat-Zr0_2_) with rms roughness below 1 nm were deposited by ion sputtering on the CPs, in order to produce reference interacting surfaces, without any nanotopographical cues. To this purpose, a Kaufman ion gun (Cyberis 40-f) was used to sputter a Zr target. The produced coating is partially oxidised in the deposition chamber; oxidation further proceeds upon exposure to ambient air.

#### Production of Poly-L-lysine –coated CPs

Also for reference purposes, CPs were coated with poly-L-lysine (PLL). PLL is a poly-amino acid routinely used to facilitate protein absorption and the attachment of cells to solid surfaces in biological applications, including our previous experiments with PC12 cells^44, 64^. For the PLL coating, the probes were incubated with a 0.1% (w/v) PLL solution (Sigma-Aldrich) at room temperature for 30 min, and washed thoroughly afterwards with milliQ water several times before the measurements.

### 2.3 Force spectroscopy experiments and data analysis

#### Force spectroscopy

The force spectroscopy experiments were performed using a Bioscope Catalyst AFM (Bruker). During the AFM measurements, the temperature of the medium was maintained at 37°C using a perfusion stage incubator and a temperature controller (Lakeshore 3301, Ohio, USA). The colloidal probes were incubated with the cell culture medium for >30 min at 37°C before the actual measurements.

The deflection sensitivity was calibrated *in situ* and non-invasively before every experiment by using the previously characterised spring constant as a reference, according to the SNAP procedure described in Ref.^41^ The standard approach, i.e. pressing the probe on a stiff surface and measuring the inverse of the slope of the force curve in the contact region, could likely cause contamination of the nt-CP surface and damage of the nanotopographical asperities. Moreover, friction-dependent issues can influence the accuracy of the determination of the deflection sensitivity by the standard contact method, when using large CPs ^83^.

Sets of raw deflection versus approaching distance curves were acquired at locations on the cells body selected by means of the optical microscope. The raw curves where converted into force versus distance curves (shortly force curves, FCs), rescaling the deflection axis by multiplication by the deflection sensitivity and the cantilever spring constant, and summing the cantilever deflection to the Z-piezo displacement axis^84^ (see Figure 3a). FCs containing 8192 points each were recorded on cells, with ramp length *l* = 8 μm, maximum load *F_max_* = 1 nN and retraction speed at *v_r_* = 16 μm/s. The pulling speed was kept relatively low to reduce hydrodynamics effects^85^.

**Figure 3.**
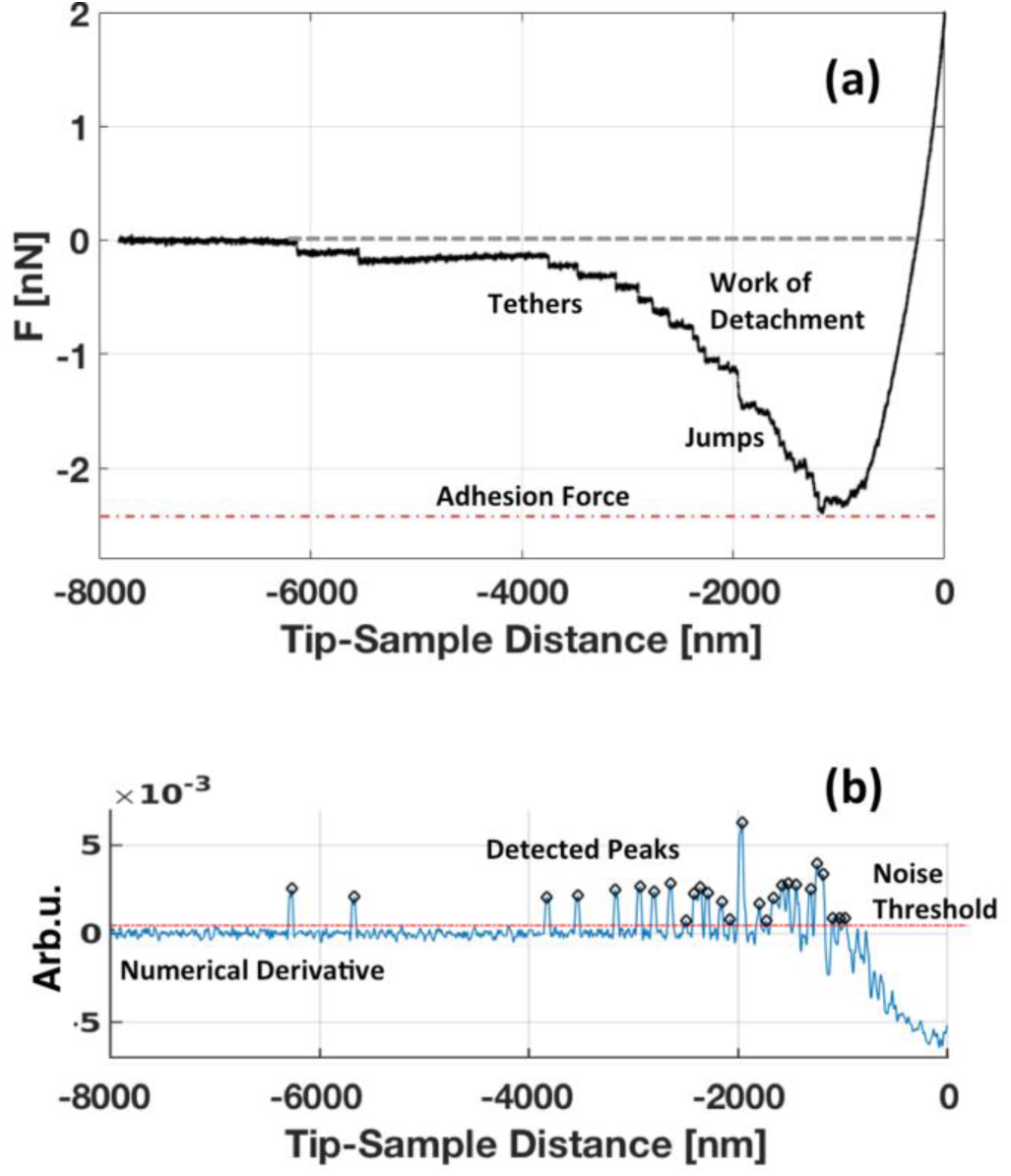
**(a)** The retraction part of a representative FC, with the two different possible unbinding events, jumps and tethers, total adhesion force and work of detachment shown. **(b)** Numerical derivative of the FC used to identify and locate the unbinding events.

To measure the early steps of cellular adhesion, we selected five contact times (*cts*): 0, 20, 60, 120, 240 s, accordingly. During the contact time, the Z-piezo position was kept constant using the Z closed-loop feedback mode. Long contact times require a very stable system; this condition is obtained by means of both an active anti-vibration base (DVIA-T45, Daeil Systems) and an acoustic enclosure for the AFM (Schaefer Italy), by controlling the environmental temperature in the laboratory, and by allowing for a long equilibration time (30 min) before starting the experiments. Figure S3 shows that the drift along the vertical direction during contact of the CP with the sample is very small (max. 20 nm in 240 s), and determines a negligible variation of the applied force.

To reduce the stress on the cells and not to alter their adhesive behaviour, a maximum applied load of 1 nN, corresponding to a pressure of the order of 10 Pa, was set during the acquisition of the FCs. For the same reason, the number of FCs collected per cell was limited (not only because of the very long acquisition time of each curve).

#### Data analysis

Data processing of the sets of curves was carried out in Matlab (Mathworks) environment using custom routines.

The values of several parameters were calculated for the analysis of the cell-probe detachment mechanisms (Figures 3a,b). In particular, we inferred the maximum cell-probe adhesion force *F_a_* and the work of detachment *W* required to separate the nt-CP from the cell. *W* is calculated as the area below the force versus distance curve. The maximum detachment force *F_a_* depends on different properties of the cell, such as overall rigidity, cortex tension, cell shape, single binding strength and spacing. Compared to *F_a_*, *W* provides a more complete, integrated information about the cell adhesion, related to the numbers, lengths and strengths of all bonds formed between the nt-CP and the cell. Moreover, we extracted the number *N_j,t_* and strength *F_j,t_* of the single unbinding events of every FC measured (for both, jumps *j*, and tethers *t*). The unbinding events must be identified in the retraction section of the FCs^86, 87^: we exploited the numerical derivative of the curves with respect to the tip-sample distance^88^ (Figure 3b) to detect the location of the events.

#### Statistics and error analysis

Mean values *ψ_mean_* and associated errors *σ_mean_* (based on the standard deviation of the mean) were calculated for each observable *ψ(F_a_, W, N_j,t_,…)*, as described in detail in the Supporting Information. These values and errors represent a population of cells in a given condition. To this purpose, first mean values and errors have been calculated for the single cells tested, then these values have been averaged and the resulting error calculated.

For each nt-CP used, 10 cells were measured (with 3 FCs for each cell). This resulted in 30 FCs per time point for each probe, and 50 cells and 150 FCs investigated for each probe in total.

### 2.4 Cell Culture and preparation for the force spectroscopy experiments

As cellular model we used neuron-like PC12 cells (i.e., in particular the PC12-Adh clone, ATCC Catalogue No. CRL-1721.1TM). These cells interact with ns-ZrO_2_ films, reacting with mechanotransductive responses to the provided nanotopographical stimuli^44, 58, 64^.

For routine cell culture (subculturing every 2-3 days), the cells were kept in an incubator (Galaxy S, RS Biotech) at 37°C and 5% CO_2_, in RPMI-1640 medium supplemented with 10% horse serum, 5% fetal bovine serum, 2 mM L-Glutamine, 10 mM HEPES, 100 units/mL penicillin, 1 mM pyruvic acid, and 100 µg/mL streptomycin (all reagents from Sigma Aldrich, if not stated otherwise).

For the force spectroscopy experiments, the cells were detached from the cell culture flasks with trypsin/EDTA solution, counted with an improved Neubauer chamber, and plated in low concentration of 4.000 cells/cm^2^ (to guarantee the presence of single separated cells) on Ø 40 mm glass-bottom dishes for cell culture (Willco Wells) the day before the experiment. Phenol-red free solutions were used for the experiments, since this molecule was found to be harmful for the AFM tip holder. Directly before the cell plating, the Ø 40 mm glass-bottom dishes were coated with PLL (incubation with a 0.1% PLL solution for 30 min at RT, followed by several washing steps with PBS) and sterilised with UV light for 10 min. After cell plating, the cells were kept overnight in the incubator to guarantee good cell attachment before the force spectroscopy experiments.

For the integrin activation, the cells were pre-incubated with manganese chloride (MnCl_2_) at a concentration of 1 mM for >10 min before the measurements (the treatment has been labelled Mn^2+^ in figures). In case of inhibition of the β1 integrin activity, the cells were pre-incubated with the inhibitory 4b4 antibody (Beckman Coulter) at a concentration of 5 µg/mL for >15 min before the measurements (the treatment has been labelled 4b4 in the figures).

## 3. RESULTS AND DISCUSSION

### 3.1 Characterisation of the nanotopographical colloidal probes

Figure 4 shows representative AFM images of a clean borosilicate glass sphere before (Figure 4a) and of the same sphere after deposition of the ns-ZrO_2_ film (Figure 4b) used for the production of a nt-CP.

**Figure 4:**
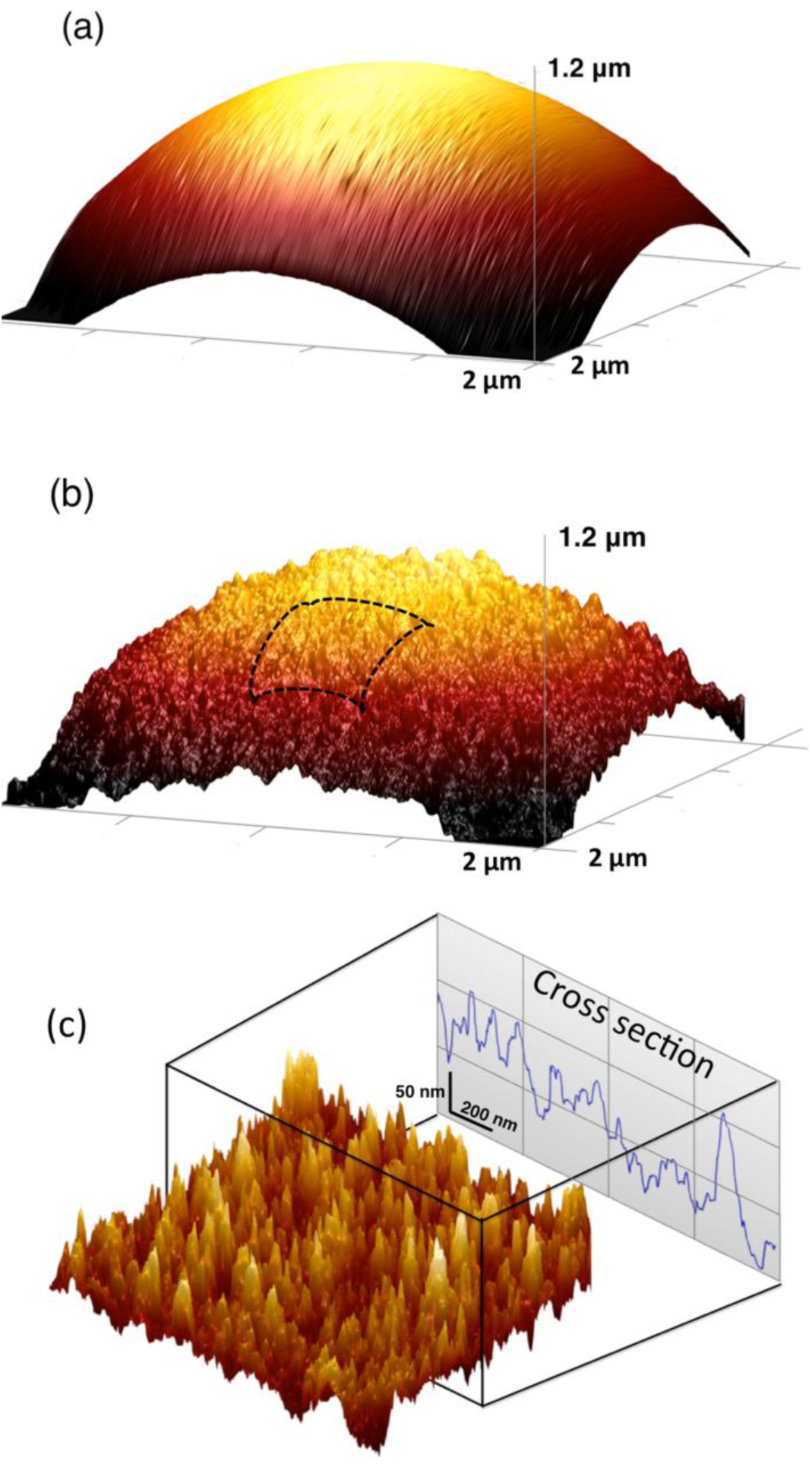
The surface morphology characterised by AFM of a borosilicate glass sphere. **(a)** Before, and **(b)** after the deposition of ns-ZrO_2_ (r_q_ = 15 nm). **(c)** A higher resolution AFM image (the baseline curvature was removed) from the nt-CP shown in Figure 4b; the cross section highlights distinct peaks and valleys at the nanoscale.

Ns-ZrO_2_ grows on the curved CP surface, as it does on conventional flat smooth substrates (see Supporting Information for details).

The structure and morphology of these cluster-assembled films result from the random stacking and aggregation of impinging nanometer-sized building blocks (the ZrO_2_ clusters) into larger and larger units. The surface profiles of nanostructured zirconia films are characterised by peaks and valleys (see Figure 4c), defining a complex random pattern of nanoscale features, whose dimensions and spatial distribution resemble those found in natural ECM topographies^5,6^. The specific surface area, the rms roughness, the average lateral dimensions of the largest morphological features (the correlation length ξ), as well as the interfacial porosity of the films typically increase with film thickness^59, 60, 89^. The interfacial open pores, delimited and defined by the surface asperities^44^, can accommodate proteins (including fibronectin) and nutrients^31, 89–92^; asperities evolve in height, area, and surface charge density^81^.

In this work, we concentrated our attention on nt-CPs with rms roughness of the ns-ZrO_2_ film *r_q_* = 15 nm. This particular value of *r_q_* was chosen because we recently demonstrated that it induces in PC12 cells mechanotransductive modulations at the level of integrin adhesion complexes and cytoskeleton (Figure 1b), as well as differentiative events, such as neuritogenesis, and a vast change in the cellular program^44, 58, 64^.

### 3.2 Cell adhesion dynamics at nanotopographical interfaces

To investigate in which way nanotopographical features influence the characteristics of cellular adhesion processes, we performed adhesion force spectroscopy on PC12 cells with four CPs, possessing different types of functionalisation:

1. ns-ZrO_2_ –coated CP, produced by SCBD, with *r_q_* = 15nm (surface: ns-ZrO_2_).
2. flat ZrO_2_ –coated CP, produced by ion gun sputtering, without nanotopographical features (surface: flat-ZrO_2_).
3. borosilicate glass CP (surface: glass).
4. poly-L-lysine –coated CP (surface: PLL).

PLL coatings are routinely used to facilitate protein absorption and cell adhesion to solid surfaces (in particular glass) in biological applications (by providing positively charged sites favouring electrostatic interactions), which represents also the canonical substrate condition for PC12 cell experimentation. In our previous work, we found that the PC12 cells formed FA and stress fibres on the PLL-coated glass and flat zirconia substrates, whereas on the ns-ZrO_2_ (*r_q_* = 15nm) substrate the adhesion sites remained at focal complex dimensions and the stress fibre formation was reduced^44^ (Figure 1a). CPs number 2, 3 and 4 were therefore used as controls.

The dependence on the contact time *ct* of the measured parameters (maximum adhesion force, work of detachment, number of unbinding events for jumps and tethers) is discussed in the next paragraphs.

#### Maximum adhesion force F_a_

Figures 5a,b show that the cells are capable to create stronger adhesion on all functionalised surfaces, compared to the untreated glass. Nevertheless, the extent of these differences compared to the reference glass surface, and the temporal evolution of the adhesion, are different on each surface.

On glass and flat zirconia, *F_a_* rapidly reaches a plateau, although on flat-ZrO_2_ the final value is higher. On ns-ZrO_2_ and PLL, *F_a_* follows a similar growing trend in the first 60 s, reaching much higher values than flat-ZrO_2_. PLL is the only surface where adhesion increases during the whole time interval, achieving its maximum values at 240 s (with the highest value of all the conditions). On the nanotopographical surface, the *F_a_* maximum is reached at 120 s, then adhesion decreases to a value similar to that on flat-ZrO_2_.

#### Work of Detachment W

The temporal evolution of the work of detachment *W* provides additional information about the level of complexity and the maturation of the cellular adhesion (Figure 5 c,d). While the trends of *W* for glass and PLL-coated glass are similar to those of *F_a_*, this is not the case for zirconia surfaces; in this case, while the evolution of *F_a_* was different for the two surfaces, the evolution of *W* is similar. Moreover, the measured work for both flat and ns-ZrO_2_ never reaches the value of the PLL-coated glass. Since the work is a force times a distance, a different trend of W compared to *F_a_* can be attributed to either different numbers of bonds (detected as unbinding events), or to different bond lengths, or both.

**Figure 5:**
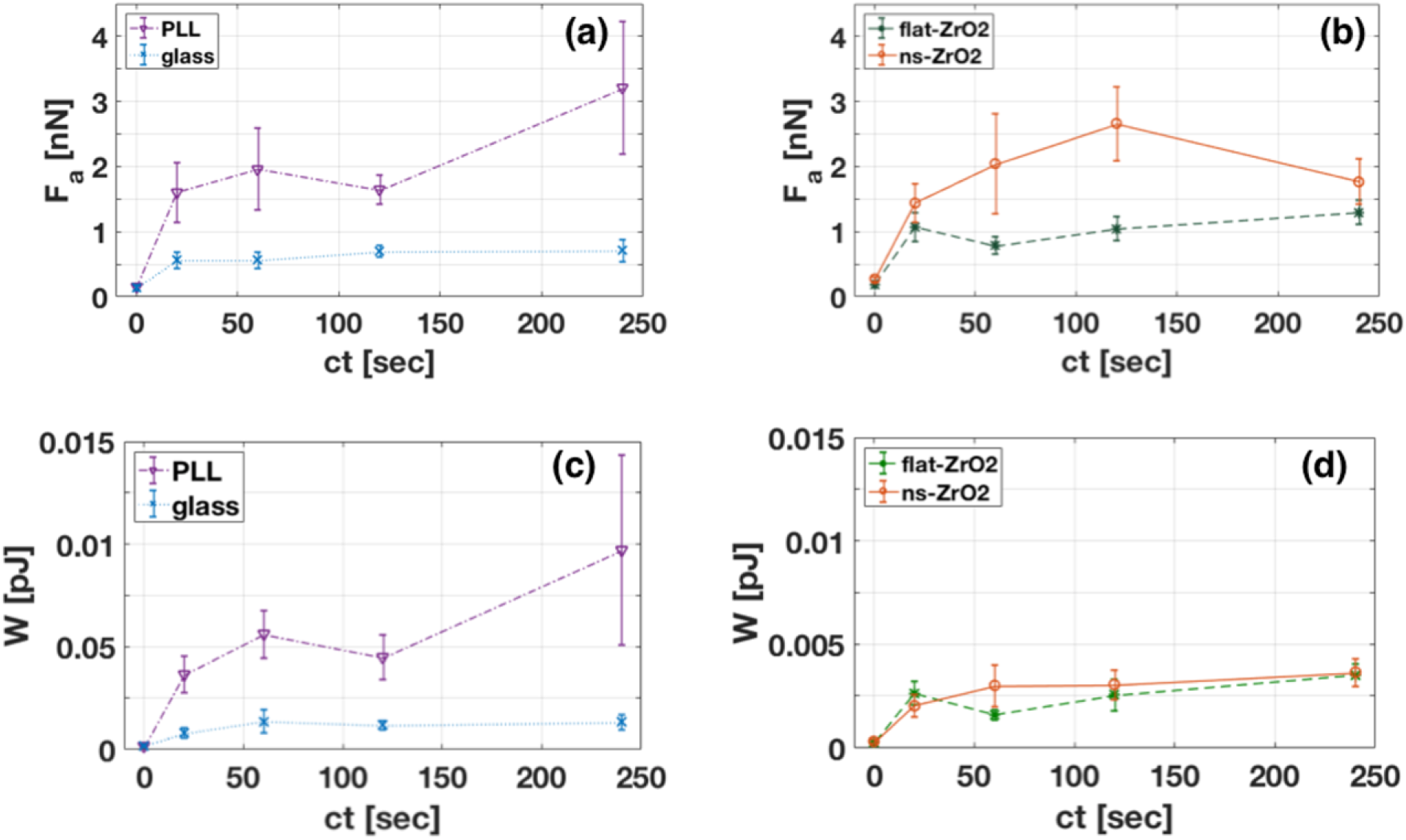
Dependence of the maximum adhesion force on the contact time for the four investigated surfaces. **(a)** On the left, the colloidal glass probes, untreated (glass, blue line) or with PLL functionalisation (PLL, violet line). **(b)** On the right, the two colloidal probes decorated with zirconia films, flat (flat-ZrO_2_, green line) or with nanotopographical features with a roughness of r_q_ = 15 nm (ns-ZrO_2_, orange line). **(c, d)** Dependence of the work W on the contact time for the same surfaces as in (a,b).

#### Mean number of unbinding events N_j_, N_t_

The mean number of detected unbinding events for cells interacting with different surfaces (Figure 6) revealed distinct differences in the temporal adhesion dynamics between PLL, ns-ZrO_2_ and flat-ZrO_2_, in particular with respect to the jump events *N_j_*, which we discuss first (Figure 6a,b).

**Figure 6.**
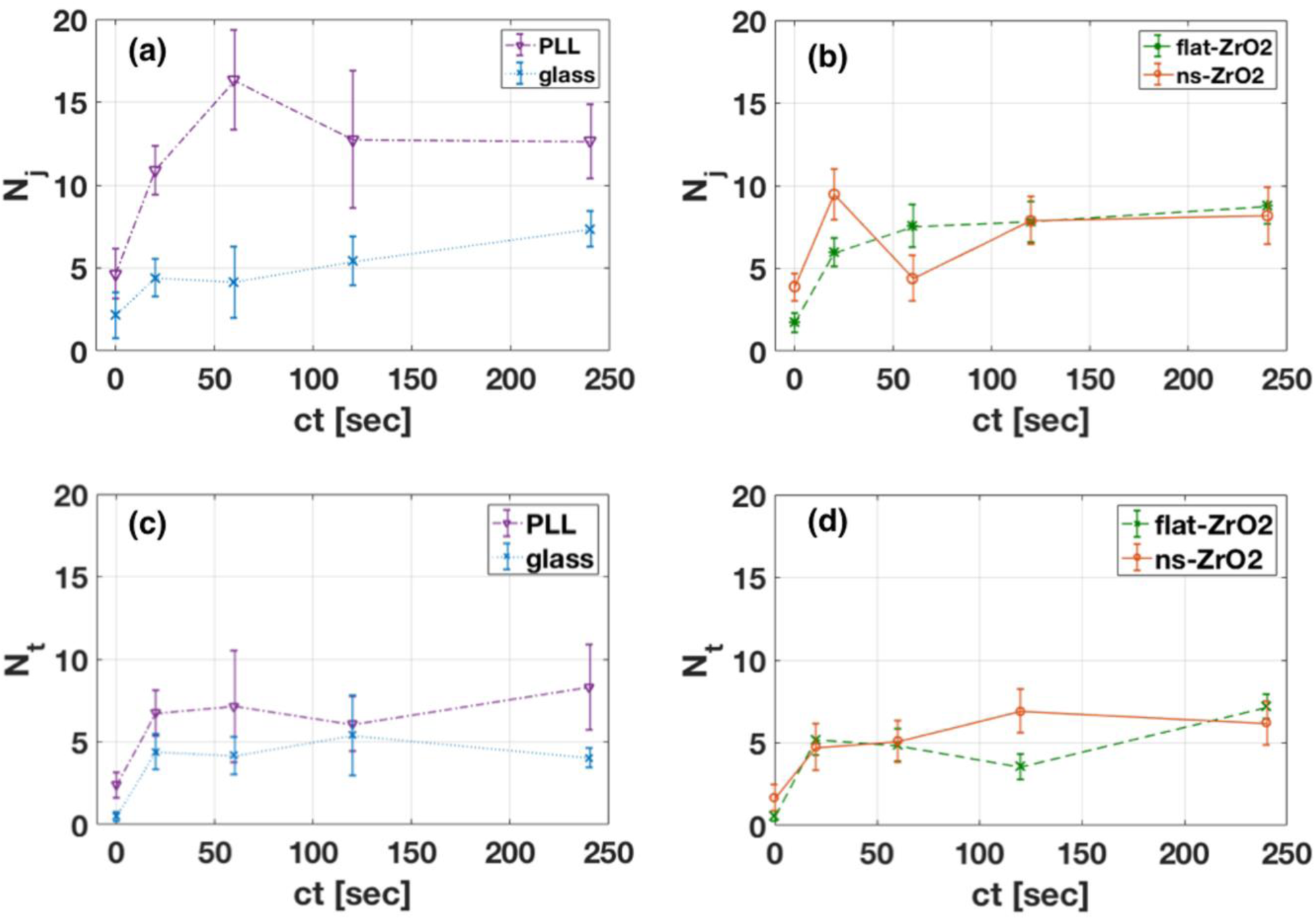
Dependence of the mean number of unbinding events on the contact time. **(a,b)** Mean number of jumps events N_j_ (see Figure 2 for details) for **(a)** the colloidal glass probes, untreated (glass-CP, blue line) or with PLL functionalisation (PLL, violet line), and for **(b)** the two colloidal probes with zirconia films, flat (flat-ZrO_2_, green line) or with nanotopographical features with a roughness of r_q_ = 15 nm (ns-ZrO_2_, orange line). **(c,d)** The same as in (a,b) for the mean number of tether events N_t_ (see Figure 2 for details).

The jump events in force curves are predominantly attributed to receptors in the membrane that are anchored to the cytoskeleton (as e.g. integrins in molecular clutches via talin)^48, 51–54, 93–95^.

PLL-coated glass (Figure 6a) and ns-ZrO_2_ (Figure 6b) show a similar progression of the *N_j_* in the first 20 s (with both reaching 10 events), after which they develop differently. The *N_j_* of PLL reaches a maximum at 60 s, and also later on the cells created significantly more jump adhesion spots on the PLL compared to flat and ns-ZrO_2_.

In the nanotopographical surface condition, *N_j_* decreases strongly (−53%) from 20 s to 60 s. Intriguingly, this drop is a recurrent and specific phenomenon that appears systematically for all investigated cells interacting with the nanotopographical surface. In the flat-ZrO_2_ condition, there is instead a progressive rise of *N_j_* in the first 120 s (reaching 8 events and being always higher than glass).

The evolution of the number of tethers *N_t_* (Figure 6c,d) instead is more similar for all four conditions.

The nature of the tethers has been poorly investigated, but they are usually associated to receptors that are not anchored to the internal actin cortex, which results in membrane extrusion from the cell reservoir^93–95^. Another hypothesis is that tether events could, at least partially, be related to the unfolding of glycocalyx sugar chains^96^. It has been demonstrated that the tethers do not respond as a catch bond^52^, and also in our experimental set-up they seem to participate in a negligible manner to the maturation of the adhesion (with a generally low contribution to *F_a_*, see Supporting Information, Figure S4), showing only minor divergent reactions towards the different surface conditions.

In the following, we will concentrate our attention mainly on the *N_j_* parameter.

The interaction time window between 20 and 120 s seems to be the most interesting for the dynamics of the adhesion spots on different surfaces, in particular when looking at the combined evolution of the different parameters (e.g., *F_a_* and *N_j_*).

PLL and ns-ZrO_2_ have a comparable development of *F_a_* from 20 to 60 s, whereas the *N_j_* evolve in a converse manner, i.e. it increases markedly for PLL, and decreases for the nanotopographical surface (Figure 6a,b). Glass and flat-ZrO_2_ instead show more moderate alterations.

In order to better investigate these dynamical phenomena, we have calculated the mean adhesion force per jump <*F_j_>* (see Supporting Information for details, Figure S4). The result is shown in Figure 7. An interesting outcome specific for the ns-ZrO_2_ surface is visible. While the average force per single jump (between 20 - 120 s) is similar for all the surfaces without nanotopographical features, i.e. glass, PLL, and flat-ZrO_2_, a sudden 3.3-fold increase of the single jump strength (from 20 s to 60 s) is evident for ns-ZrO_2_ (for flat-ZrO_2_ it actually drops by 50% in the same time frame). At regime, after 240 s, the force <*F_j_>* converges down to the value of the other surfaces, nevertheless it remains significantly higher compared to glass and flat-ZrO_2_ for most of the time.

**Figure 7.**
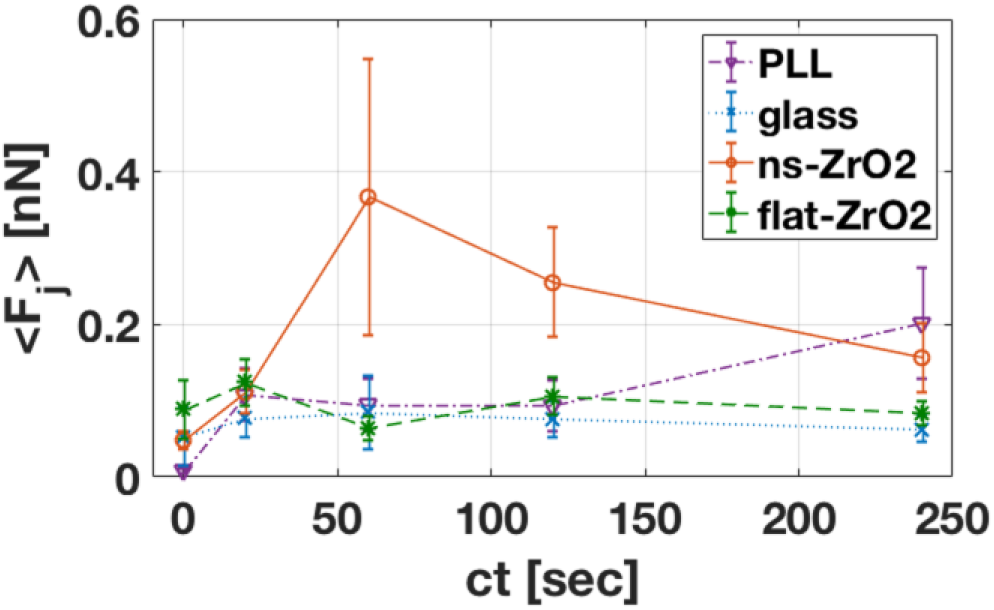
Evolution of the mean jump force versus the contact time for the different surfaces (glass: blue line; PLL: violet line; flat-ZrO_2_, green line; ns-ZrO_2_, orange line).

Interestingly, these divergent dynamics happen in the critical time window for molecular clutch reinforcement, the initiation of integrin clustering and nascent adhesion growth^52, 97^.

### 3.3 Adhesion dynamics to the nanotopographical probe are influenced by the availability of activated integrin and depend, at least partially, on β1 integrin

Due to our previous data on the effects of the nanotopography on the configuration of integrin adhesion sites^44^ and the interesting chronology of the jump adhesion events and force development observed for the ns-ZrOsurface, we tested in which way these dynamics towards the nanotopography depend on the integrin activity.

In a first step, we examined how an excess of integrin activation would affect the early adhesion dynamics towards the nanotopographical surface. To this purpose, we activated the integrins with Mn^2+^. This treatment kept the *N_j_* from dropping after 60 s, as it was observed on the ns-ZrO_2_ surface without Mn^2+^ activation (Figure 8a).

**Figure 8.**
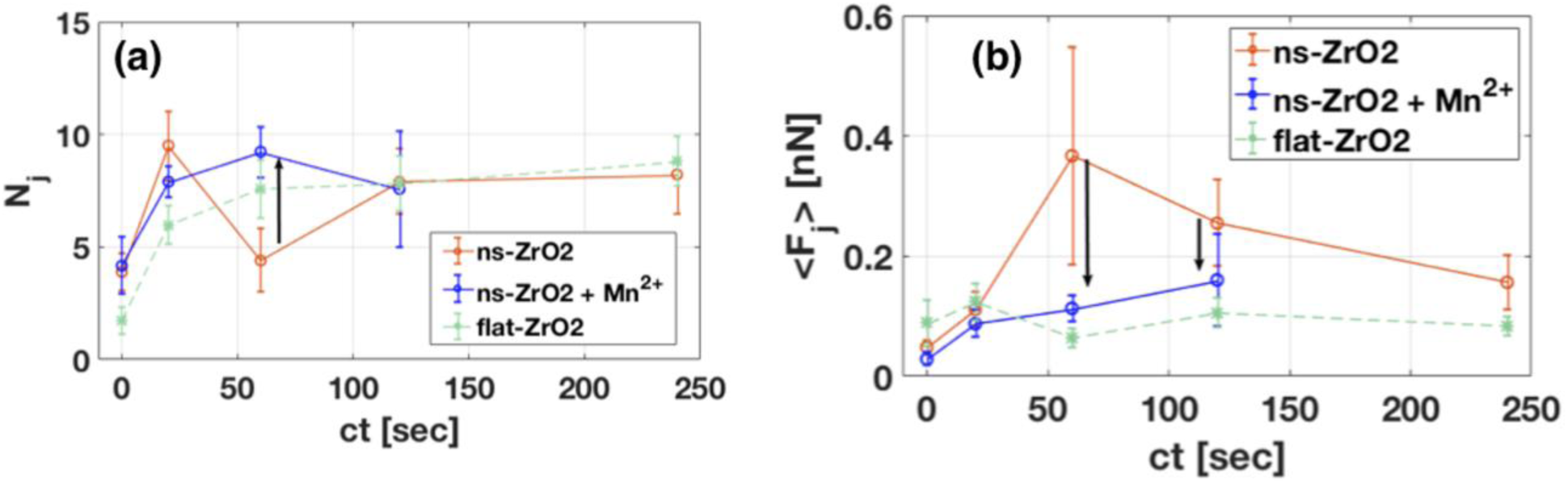
**(a)** Mean number of jumps N_j_ detected on ns-ZrO_2_ surface before (orange) and after (blue) Mn^2+^ treatment versus the contact time. **(b)** Mean jump force <F_j_> measured on ns-ZrO_2_ before (orange) and after (blue) Mn^2+^ treatment versus the contact time. The flat-ZrO_2_ cases (green, reproduced from Figure 7) are also shown for sake of comparison. The black arrows highlight the effect of the Mn^2+^ treatment on both N_j_ and <F_j_> in the ns-ZrO_2_ condition.

This impact of the abundant availability of activated integrins might be explained by two effects (or a combination of both). In nascent adhesions, Mn^2+^-induced integrin activation is known to increase the density within integrin clusters^98^; moreover, unligated activated integrins could favour the bridging between separated, but adjacent adhesion sites^99^. In any case, the force loading can be distributed over more active integrins, confirmed by the decrease of mean force per jump event at 60 s due to the Mn^2+^ treatment (Figure 8b), which stabilises the adhesion sites.

Furthermore, we treated the cells with an allosteric inhibitory antibody against β1 integrin (4b4), to see how this would impact on *N_j_* at later stages, when integrin clustering strongly manifests (120 and 240 s). As shown in Figure 9, in the presence of the 4b4, N*_j_* decreased by 42% at 120 s, and by 60% at 240 s. Since β1 integrin represents the most common, but not the only, β integrin subunit, this result demonstrates that the recorded jump interaction events depend, at least partially, on the activation of β1 integrin subunit-containing integrin receptors. This outcome is in line with the involvement of the (β1) integrin activation and signalling in the nanotopography-sensitive modulations in PC12 cell mechanotransduction and differentiative behaviour (in particular neuritogenesis) we have previously reported^44, 64^.

**Figure 9.**
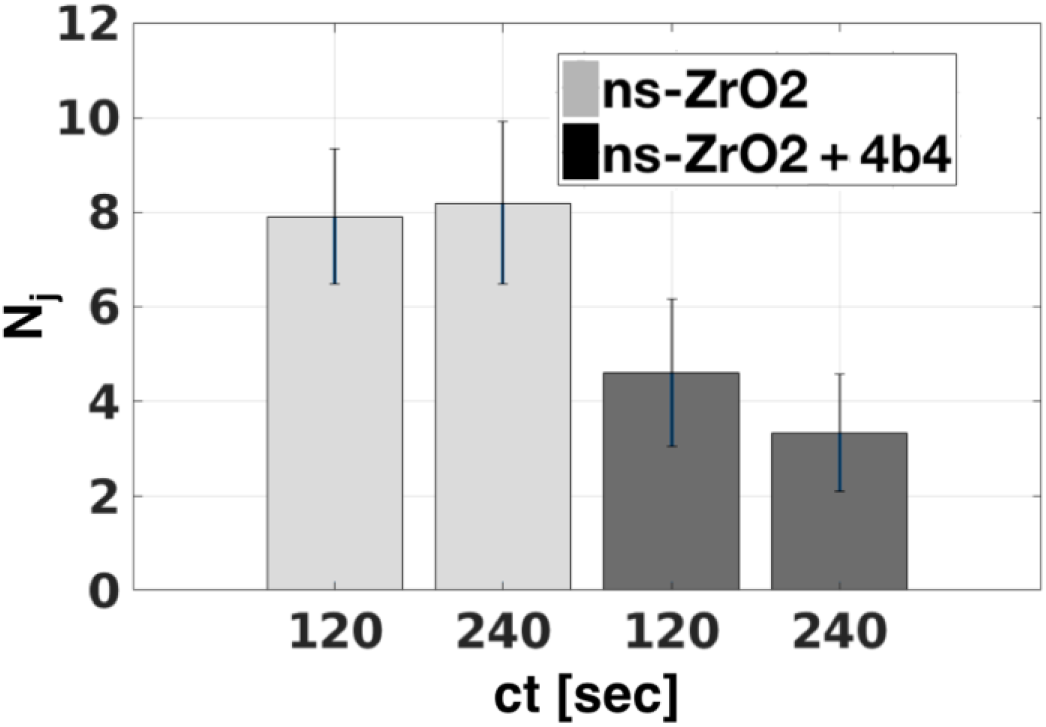
Mean number of jumps N_j_ in the ns-ZrO_2_ condition before (grey) and after (dark grey) the 4b4 antibody treatment (5µg/mL), for ct = 120 s and ct = 240 s.

Altogether, these results indicate that the availability of activated integrins seems to be an influential regulatory factor for spatial sensing of adhesion sites at the nanoscale.

### 3.4 Evolution of jump force distribution over time

For a better understanding of the dynamics of the adhesion force development on the different surfaces, we analysed in greater detail the distribution of the measured strengths of the single jumps at different contact time *ct*.

At *ct* = 0 s, the distributions of jump forces are similar for all surfaces, with a leading peak detected around 40 pN for zirconia surfaces and 20 pN for glass and PLL (see Table 1). The immediate appearance of a force peak in this range is consistent with the recently shown very fast integrin adhesion response (in that case for α5β1 integrin/fibronectin binding)^50^. The range is also compatible with the reported peak tension of single integrin-ligand bonds during initial adhesion^51, 100–104^ and furthermore, it coincides with the force thresholds for the extension of the talin rod, vinculin recruitment and molecular clutch reinforcement^105^.

With the increase of *ct*, we observed that on glass and PLL the mean jump force remains almost constant, with the appearance of a minor peak at higher forces around 120 pN (Figure 10a,c). The distribution of flat-ZrO_2_ (Figure 10b) is similar, although the peaks are broader and there are more counts at higher forces.

**Figure 10.**
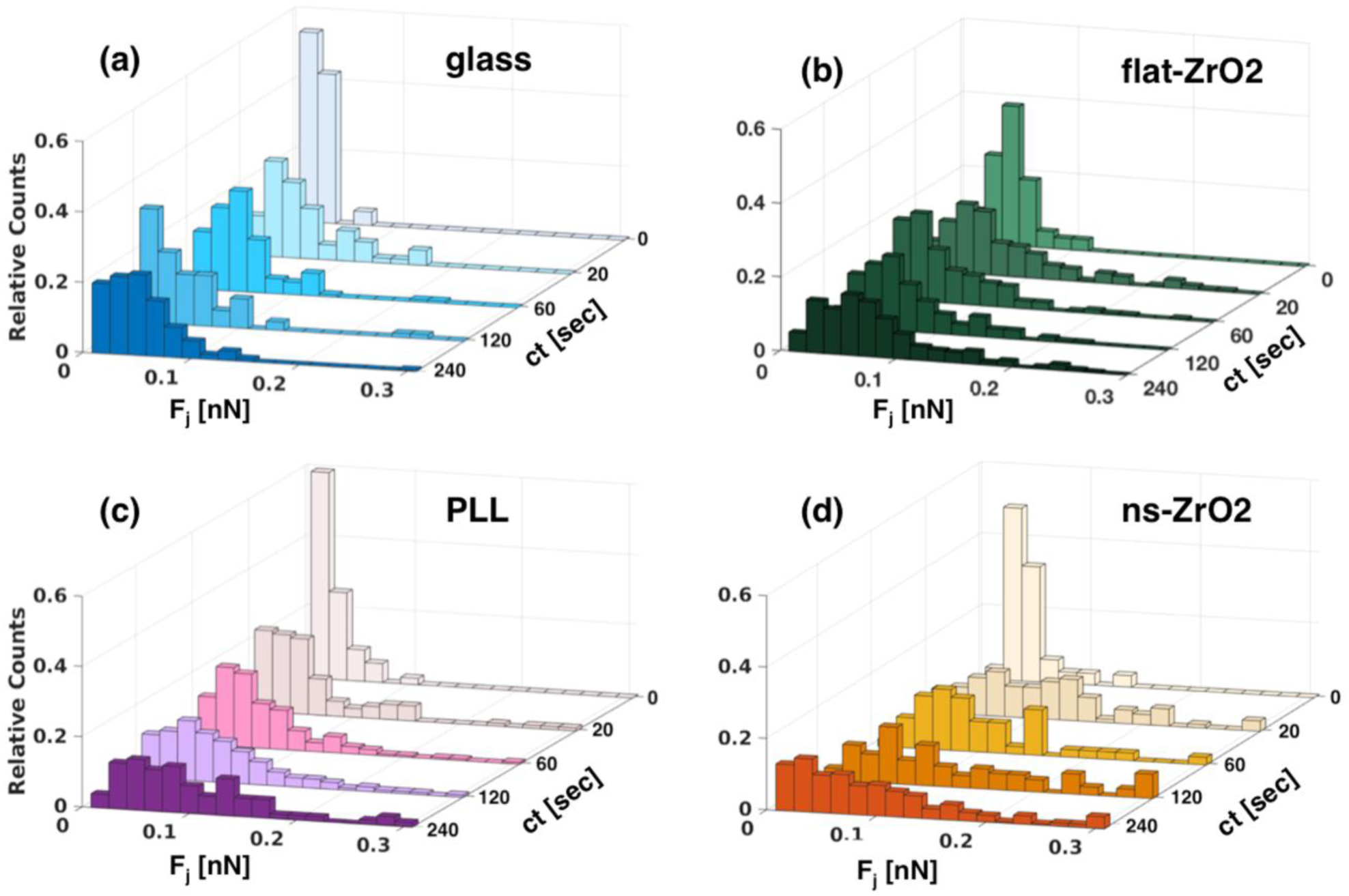
Evolution of the jump force distribution with contact time for **(a)** glass (blue bars), **(b)** PLL (violet bars), **(c)** flat-ZrO_2_ (green bars) and **(d)** ns-ZrO_2_ (orange bars). The relative counts are the abundances of unbinding events (in each bar) normalised to total number of unbinding events

**Table 1.** Most probable jump force at ct = 0s.

| Surfaces | Most probable jump force at $ct = 0$ (pN) |
| --- | --- |
| flat-ZrO<sub>2</sub> | 42±15 |
| ns-ZrO<sub>2</sub> | 36±13 |
| glass | 21±10 |
| PLL | 19±17 |

On ns-ZrO_2_ (Figure 10d) the distribution of forces, already after 20 s, shifts towards higher forces, even above 200 pN. High force bonds (>100 pN) appear almost with the same frequency (∼45%) as the weaker ones (<100 pN) at every time point (0 s excluded).

Regarding the adhesion force *F_a_* (Figure 5b), this increased strength of the single jumps actually compensates for the lower number of adhesion spots compared to the PLL-coated glass (in particular at *ct* = 60 s, see Figure 6a,b). In other words, on the nanotopographical surfaces the adhesion spots are exposed to higher forces. The decrease of high-force unbinding events on ns-ZrO_2_ observed at *ct* = 240 s, with respect to earlier *cts*, is compatible with the general decrease of the *F_a_* (Figure 5b).

We point out how the increased occurrence of higher-force events (Figure 7 and 10) happens simultaneously with the drop of *N_j_* (Figure 6a), which depend on the availability of activated integrins (Figure 8), in the ns-ZrO_2_ condition. This is peculiar, in particular because it happens during the critical time window for nascent adhesion growth and integrin clustering^97^. This temporal course of the nanotopography-specific adhesion dynamics could indicate that too small and/or too separated adhesion sites (critical thresholds for integrin clustering have been determined to be ≥60-70 nm) disintegrate, due to exposure to excessively high forces. Instead, when the conditions are suitable, bigger clusters of integrin form and succeed to reinforce and mature^55, 56, 67, 106, 107^ (Figure 11). These clusters might need to exceed at least the size of the minimal adhesion unit of a few integrins in sufficiently close, i.e. tens of nanometres, vicinity^99, 108^; which is consistent with the appearance of the second peak at 120 pN.

**Figure 11.**
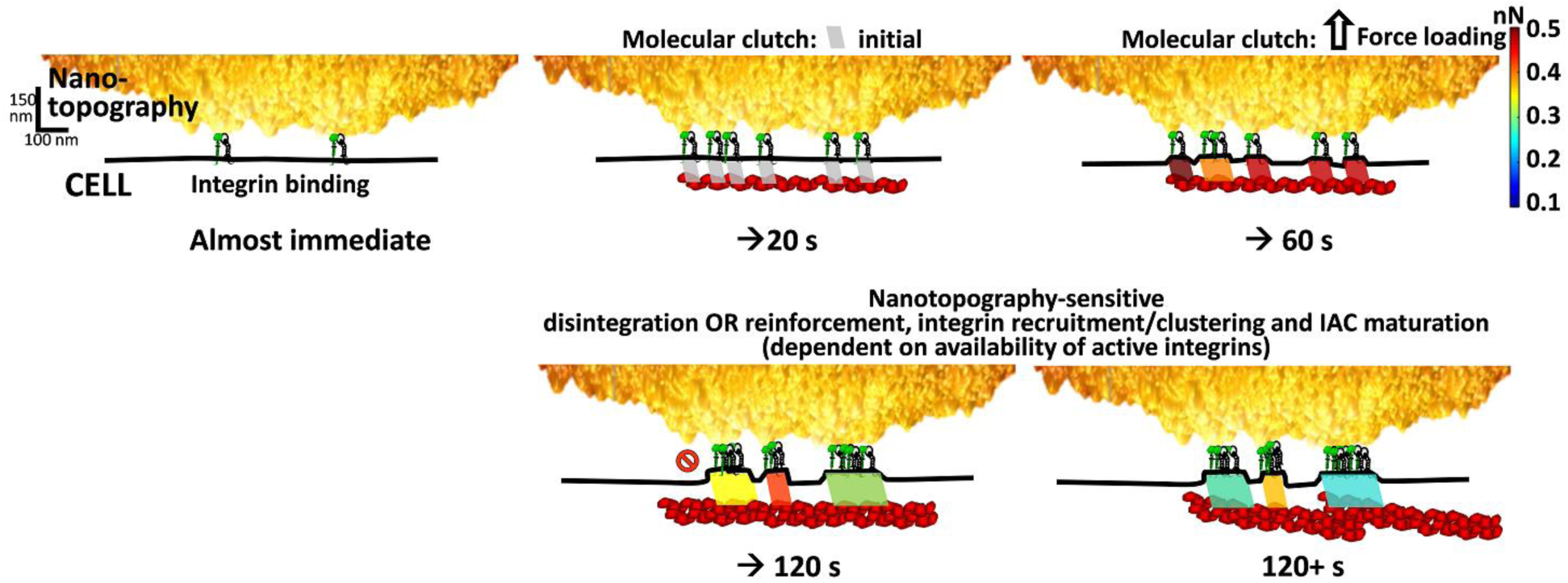
This graphics is a visual representation of our actual experimental data integrated into a potential mechanism of nanotopography-sensitive force loading dynamics (left to right, and row-wise). The colours of the parallelogram icons (symbolising the engagement between the integrins and the actomyosin-generated forces, i.e. the molecular clutches) code for the force intensities measured by means of our approach at the different time points (see the bar on the right, compare with Fig. 7). The scheme illustrates the integrin recruitment/clustering, force distribution, molecular clutch reinforcement and integrin adhesion complex (IAC) maturation where the nanotopographical conditions are suitable, or disintegration of the initial structure due to excessive force exposure (because integrin recruitment and force distribution is insufficient) where the nanotopographical conditions are not suitable. The experiments with Mn^2+^ treatment furthermore showed that these dynamics are dependent on the availability of active integrins (see Fig. 8). The length scales on the left refers to the morphology of the nt-CP only; the other graphical objects (such as the integrins) are not in scale and also their number is symbolic to visualise the development.

These results are compatible with the dimensions of the nanotopographical features and the confinement action at the nanoscale, provided by the small contact area offered by the asperities of the nanostructures and the distance between them.

We have demonstrated previously that the PC12 cells are in contact only with the apical part of the nanotopographical asperities, which restricts the size of the adhesions sites to the nanometric level, and inhibits the maturation of bigger adhesion structures on a larger scale (see Figure 1a). Analyses of TEM images (an example in Figure 1a) showed that the contact areas between the cells and the asperities of the nanotopography (with a *r*q = 15 nm) have an average width of 53.2 ±48.0 nm (median: 40.4 ±21.6 nm). Moreover, the distances (mean: 99.1 ±101.4 nm, median: 60.4 ±30.4 nm) between the asperities of the disordered nanotopography effectively oscillate around the critical ligand spacing threshold (≥60-70 nm)^44^. The results reported in this work are consistent with these previous observations, and confirm, at the level of single binding events and at the pN scale, the crucial role of force loading in the molecular clutches in the nascent adhesions to accomplish nanometric spatial sensing of adhesion sites, reported recently by Oria et al.^19^.

## 4. CONCLUSIONS

Integrin-mediated mechanosensing and mechanotransduction is regulated by biophysical properties of the cellular microenvironment, such as e.g. the nanotopography of the ECM, as it influences the spatial and geometrical organisation of potential cellular adhesion sites^7–10, 29^. However, a precise understanding of the spatiotemporal dynamics and molecular actions in the cell/microenvironment interface during adhesion is still elusive, because these events take place at the nanometre scale and involve pN range forces, whose exploration requires sophisticated methodologies. Moreover, an accurate control over the nanotopographical features of the microenvironment is essential, in order to systematically investigate and precisely assess the influence of the different nanotopographical motifs on the mechanotransductive process.

In this framework, we were able to study and quantify the impact of microenvironmental nanotopography on early cellular adhesion events by means of adhesion force spectroscopy based on novel colloidal probes mimicking the nanotopography of natural ECMs. Our approach merges the sensitivity of AFM-based force spectroscopy^36, 48^ with the possibility of controlling the nanotopographical features of the ECM-mimicking substrate provided by Supersonic Cluster Beam Deposition^58^.

Thanks to our innovative approach, we could detect nanotopography-specific modulations of the molecular force loading dynamics and integrin clustering at the level of single binding events, in the critical time window of nascent adhesion formation. Following this approach, we found that the availability of activated integrins is a critical regulatory factor for these nanotopography-dependent dynamics.

Our results are in agreement with the reported importance of force loading^19^ for cellular spatial sensing of the microenvironment, and integrin nanocluster bridging between adjacent (tens of nm) adhesion arrays^99^.

In the future, we plan to exploit the fine control of the surface topography of the nanostructured colloidal probes to dissect in more detail the precise role of morphological properties (such as, e.g., roughness, asperities diameter/distance, or correlation length) in spatiotemporal cell adhesion dynamics, in particular focusing the attention on the clustering of integrin adhesion complexes^44^. Nanotopographical colloidal probes could also be used to examine how aberrations in components of the mechanotransductive machinery alter the force loading dynamics of cellular adhesion.

Additional features of nt-CPs make them suitable for the investigation of nanoscale phenomena of nanobiotechnological interest. Due to their large area, nt-CPs allows to optically probe the interaction interface, which is particular useful when using fluorescence microscopy and suitable staining of the actin cytoskeleton to image the focal adhesion spots. Moreover, nt-CPs could be used as nanotopographical templates for carrying out further functionalisations, for example by grafting to the corrugated surface biochemical moieties relevant to mechanotransduction, such as ligands, ECM motifs, etc. Apart from the cell biology framework, we think these probes have a good potential if used in physics of matter experiments, e.g. in the context of nanofriction^109^, DLVO theory^81, 110^ or adhesive contact mechanics at disordered, nanostructured interfaces^111, 112^.

## Acknowledgements

We acknowledge the support of the European Union’s Horizon 2020 research and innovation programme under the Marie Skłodowska-Curie grant agreement No 812772, project Phys2BioMed, and under the FET Open grant agreement No. 801126, project EDIT. PM and CS acknowledge support from the European Union FP7-NMP-2013-LARGE-7 “FutureNanoNeeds” programme. We thank Francesca Borghi for support in the characterisation of CPs, Paolo Piseri for useful suggestions for the deposition of thin films by ion beam sputtering, and Mirko D’Urso for support in cell culture. We thank Dr. Stefano Marchesi and Dr. Stefania Marcotti for critical reading of the manuscript.

## Author contributions

Conceptualisation: CS, AP; methodology - probe fabrication and characterisation: MC, AP, CP; methodology - cell culture and preparation: TD, CS; methodology - AFM spectroscopy: MC, TD, AP; data curation and analysis: MC, CS, AP; original draft writing: MC, CS, AP; draft reviewing and editing: MC, CS, AP, CL, PM; supervision: CL, PM, CS, AP; resources, funding and project administration: CL, PM, AP. Author contributions were allocated adopting the terminology of CRediT - Contributor Roles Taxonomy.

## SUPPORTING INFORMATION

### Scaling of the surface roughness of ns-ZrO_2_ films

To characterise the growth mechanism of ns-ZrO_2_ films on colloidal probes (CPs), we dispersed the glass spheres on a flat glass microscopy slide, and deposited ns-ZrO_2_ films varying the deposition times. We then imaged the coated CP surfaces by AFM in Tapping Mode (probe model: NCHV, Bruker), with relative scan speed of the tip v_scan_= 2μm/s, and we measured the rms roughness *r_q_*. The film thickness *h* was measured on the flat glass surface, in a region close to a sphere, by imaging the ns-ZrO_2_ film across a sharp step produced by masking the substrate during deposition. By applying a linear regression on a loglog scale to the *r_q_* versus *h* curve (see Figure S1), the value of the growth exponent *b* can be determined as the slope of the curve, according to the equation *r*_*q*_∼ *h*^*b*^.

According to previous results^1–3^, the *b* parameter of the cluster-assembled ZrO_2_ thin films grown on flat substrates is *b* = 0.368 ± 0.001 on silicon, or *b* = 0.31 ± 0.09 on glass coated with a monomolecular PAcrAm-*g*-(PMOXA, NH_2_, Si) layer^4^. These values are compatible with the prediction of the ballistic deposition growth model (*b* = 0.32 - 0.25), which assumes that clusters impinge with a direction perpendicular to the plane of the substrate, and they do not diffuse significantly upon landing^5–7^. Higher values can be found when the impinging particles possess a distribution of size and different sticking probabilities.

On nt-CPs, we found *b* = 0.314 ± 0.017, in agreement with the value measured on flat substrates. Therefore, the curvature of the nt-CPs did not influence the growth exponent.

### Mechanical stability of ns-ZrO_2_ films

To test the mechanical stability of the nanostructured coating we used a stiff AFM tapping mode probe (force constant K = 50. 4 N/m) to apply high forces on the thin film, in order to record at witch forces rupture events between ZrO2 nanoparticles take place.

An example of the force curve (FC) with the rupture events detected is shown in Figure S2a, while the rupture forces measured are represented in Figure S2b. Rupture forces are clustered around specific values and it is possible that the higher forces events represent cascade rupture events, where groups of nanoparticles simultaneously detach, while the lower forces represent the tip slipping across a nanoparticle, or small nanoparticles detaching from low-attachment points. The lowest rupture force detected is around F∼ 70 nN, that is almost 25 times larger than the highest force measured during the force spectroscopy experiment.

Furthermore, we scanned in Tapping Mode the surface of the contact region of an nt-CP after a whole day of force spectroscopy experiments. The image obtained is shown in Figure S2c, after subtraction of the spherical curvature in order to highlight the morphological details at the nanoscale. The granularity of the surface due to the presence of the ns-ZrO2 thin film is clearly evident.

### Statistics and error analysis

For each observable *ψ _FCs_* extracted by each force curve (FC) a mean value *ψ_cell_* was evaluated for each cell. The error *σ*_*cell*_ associated to ψ_cell_ was obtained by summing in quadrature the standard deviation of the mean *σ_ψ_* of the results coming from the single FCs and an estimated instrumental error *σ_instrum_* (*σ_instrum_ / ψ* = 3%)^8^:

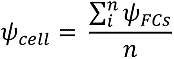

where *n* is the number of force curves per each cell,

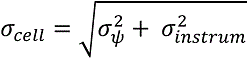

The final mean value *ψ_mean_* representative of the cell population behaviour in a given condition was evaluated as:

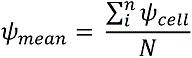

where *N* is the number of cells investigated for the given condition.

The final error *σ_mean_* associated to *ψ_mean_* was calculated by summing in quadrature the propagated error of the mean *σ_s_* and the standard deviation of the mean of the singles cell values *ψ_cell_*:

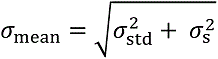

where

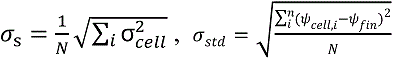

### Force curves during contact

To test the stability of the Z-piezo upon contact of the CP with the sample in close-loop mode, we recorded the cantilever deflection as a function of the contact time *t*, up to *t* = 240s.

The time-dependent deflection is shown in Figure S3. On a stiff substrate, a smooth drift is observed (∼10nm in 240 s), corresponding to a maximum variation of the applied force of approximately 0.5 nN after 240s. On the cell surface, a similar drift is observed superimposed to small and slow (tens of nm in tens of seconds) fluctuations, due to the adjustment of the cell below the probe, probably accompanied by internal reorganisation of the cytoskeleton^9^.

### Dependence of the adhesion force on the contact times, with the contribution of the tethers

The evaluation of the tethers contribution to the total adhesion force *F_a_* was evaluated as follow: for each contact time, the mean value of the tethers unbinding force has been evaluated and multiplied by the mean number of tethers *N_t_* per FC, to obtain the tether background adhesion value.

The contribution of the tethers then has been subtracted from the adhesion force *F_a_*, in order to calculate the contribution of the jumps. The obtained total jump adhesion force was eventually divided by the mean number of jumps *N_j_*, in order to calculate the mean adhesion force per jump presented in Figures 7,8 of the main text.

Representative retraction force curves at contact time *ct* = 60 s

We show representative of FCs of the adhesive behaviour of the cells on the different surfaces for contact time *ct* = 60s with jump events highlighted by black arrows.

It is possible to observe the differences in the adhesion force *Fa*, work of detachment *W*, and number of jumps *Nj*, respectively. In particular, it can be seen how the adhesion on the ns-ZrO2 surface reaches values compatible to the adhesion on the PLL, even in the presence of much less detachment events.

---

**Figure S1.**
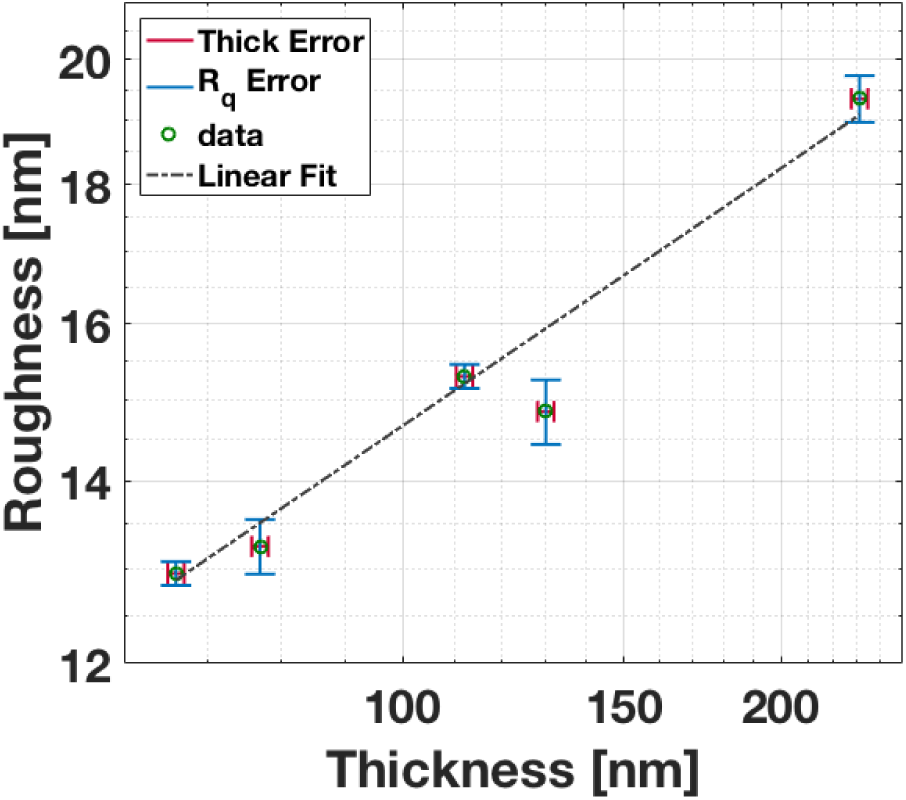
Scaling of the rms roughness r_q_ of ns-ZrO_2_ film (i.e. roughness) on the nt-CPs.

**Figure S2.**
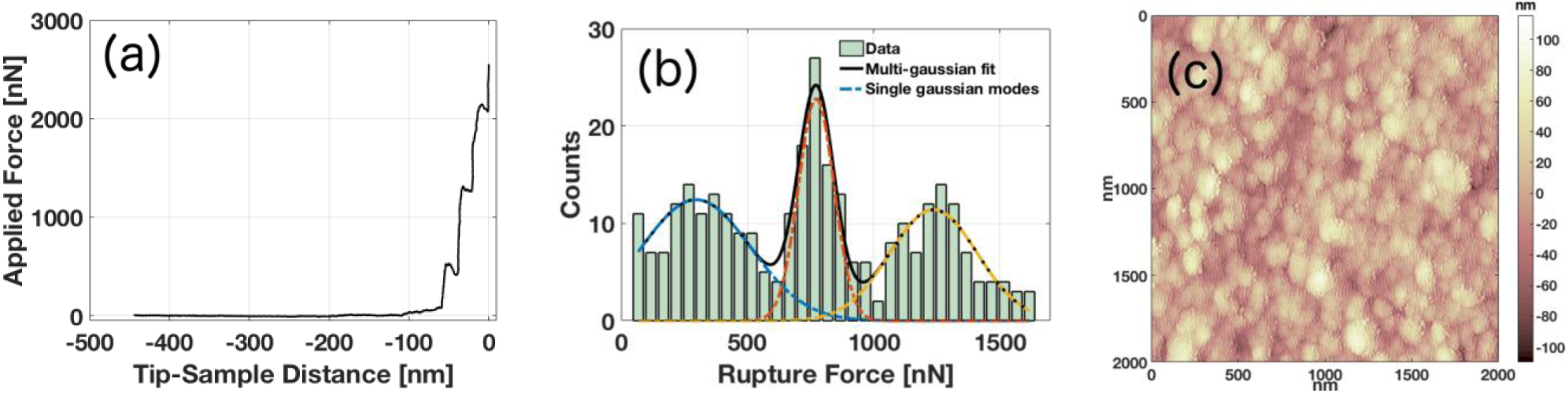
(a) Representative FC with three rupture events detected; (b) distribution of the rupture event forces from all FCs, with multi-gaussian fit highlighted; (c) scan of the contact surface of an nt-CP after force spectroscopy experiment.

**Figure S3.**
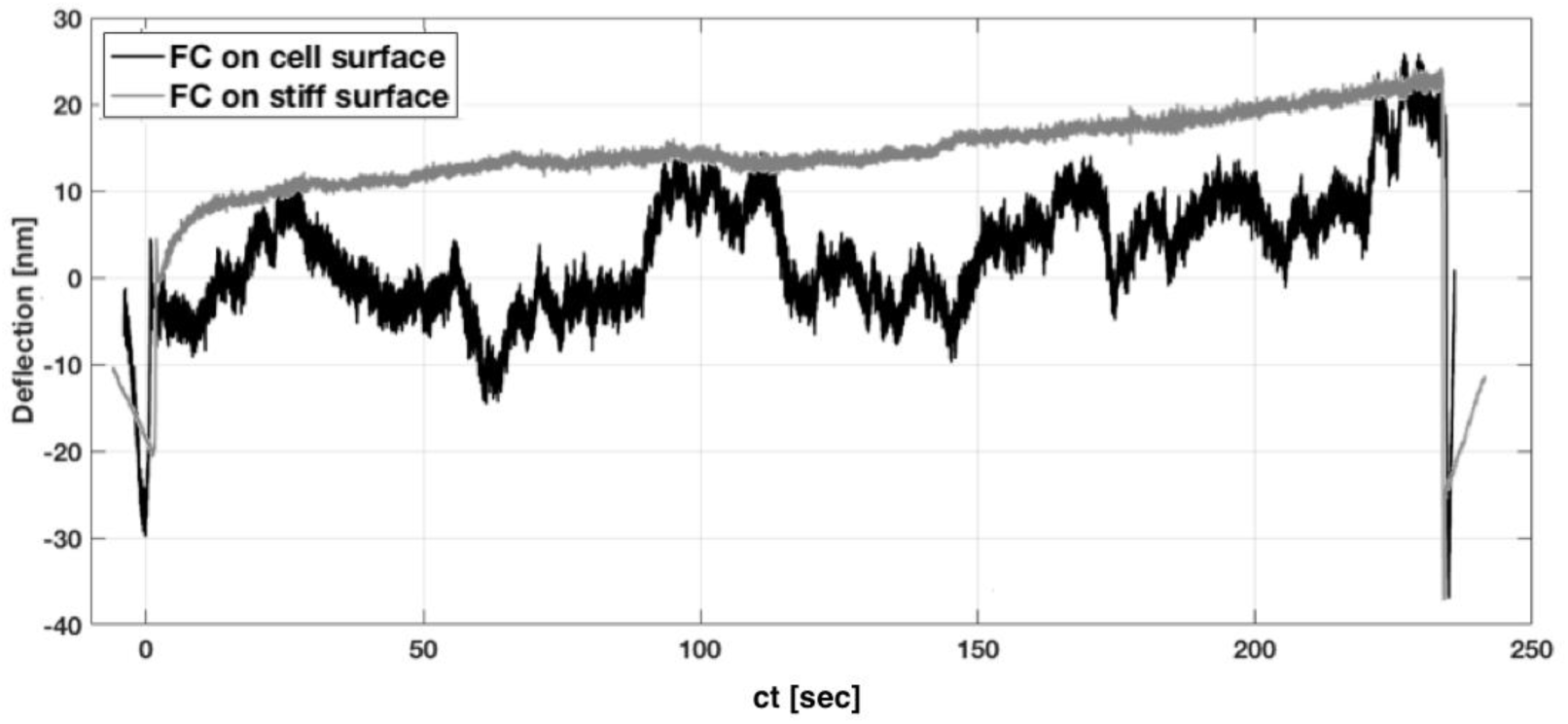
Representative deflection versus contact time curves.

**Figure S4.**
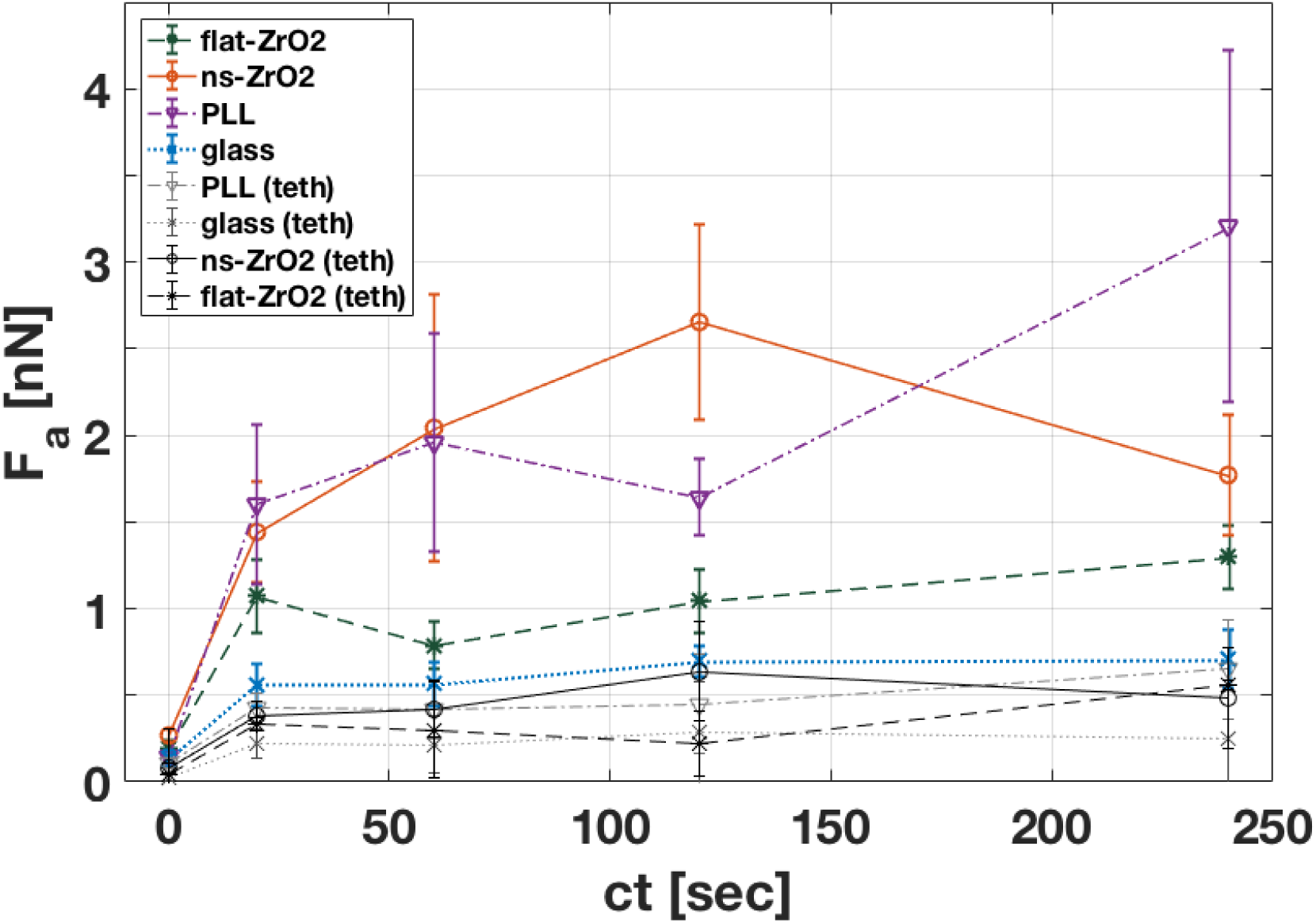
Dependence of the adhesion force on the contact times as in Figure 5; in addition, the contribution of the tethers is shown.

**Figure S5.**
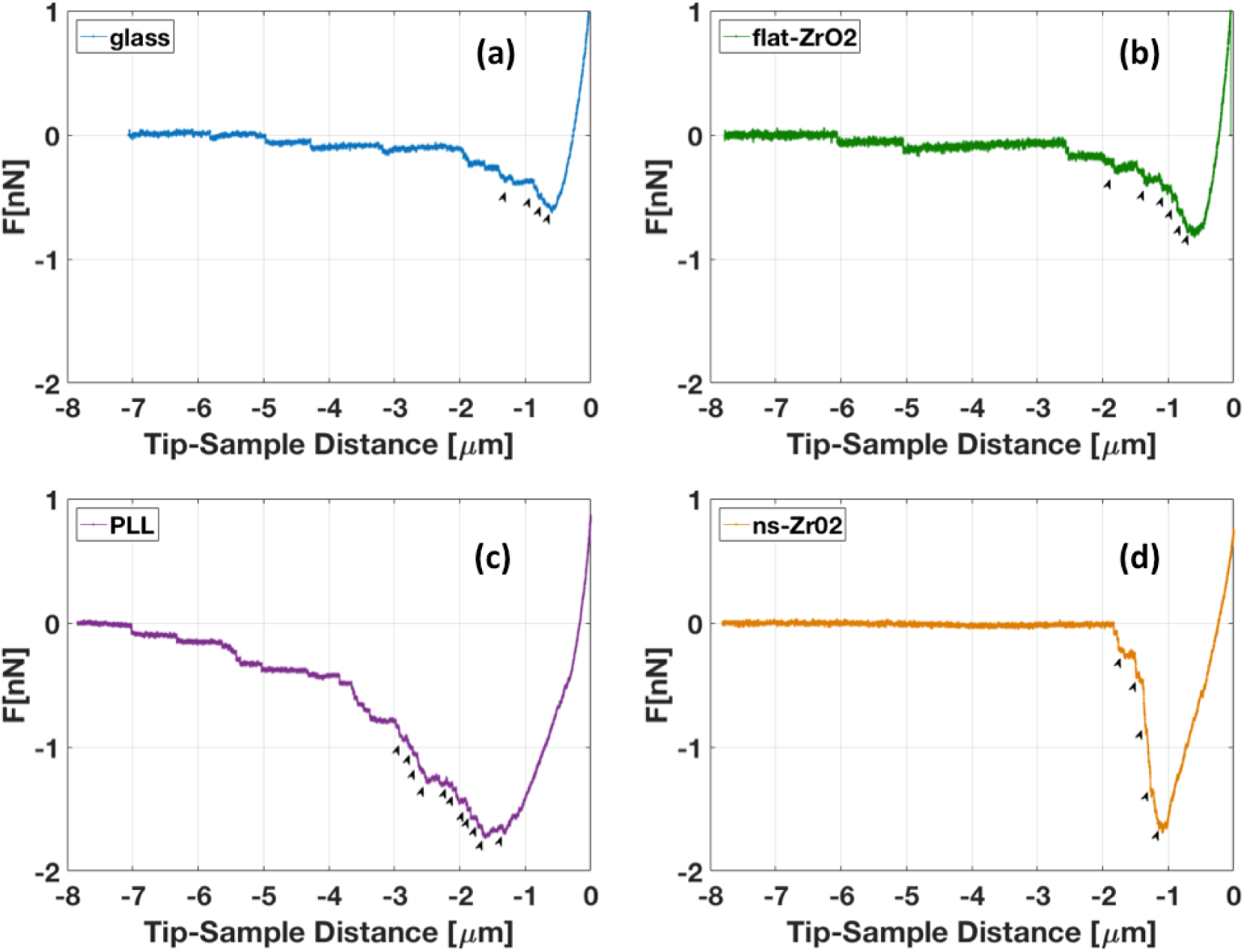
Representative FCs at ct=60 s for each condition: (a) glass, (b) flat-ZrO_2,_ (c) PLL, (d) ns-ZrO_2_.

